# Dynein acts to cluster glutamate receptors and traffic the PIP5 kinase, Skittles, to regulate postsynaptic membrane organization at the neuromuscular junction

**DOI:** 10.1101/2021.09.27.462070

**Authors:** Amanda L. Neisch, Thomas Pengo, Adam W. Avery, Min-Gang Li, Thomas S. Hays

**Affiliations:** Department of Genetics, Cell Biology, and Development, University of Minnesota, Minneapolis MN; University of Minnesota Informatics Institute, University of Minnesota, Minneapolis, MN; Department of Chemistry, Oakland University, Rochester, MI

## Abstract

Cytoplasmic dynein is essential in motoneurons for retrograde cargo transport that sustains neuronal connectivity. Little, however, is known about dynein’s function on the postsynaptic side of the circuit. Here we report distinct postsynaptic roles for dynein at neuromuscular junctions (NMJs). Intriguingly, we show that dynein punctae accumulate postsynaptically at glutamatergic synaptic terminals. Moreover, Skittles, a phosphatidylinositol 4-phosphate 5-kinase that produces PI(4,5)P_2_ to organize the spectrin cytoskeleton, also localizes specifically to glutamatergic synaptic terminals. Depletion of postsynaptic dynein disrupts the accumulation of Skittles, PI(4,5)P_2_ phospholipid, and organization of the spectrin cytoskeleton at the postsynaptic membrane. Coincidental with dynein depletion, we observe an increase in the clusters size of ionotropic glutamate receptor (iGluR), and an increase in the amplitude and frequency of mEJPs. However, PI(4,5)P_2_ levels do not affect iGluR clustering and dynein does not affect the protein levels of iGluR subunits at the NMJ, suggesting a separate, transport independent function for dynein in iGluR cluster organization. As dynein punctae closely associate with iGluR clusters, we propose that dynein physically tethers iGluR clusters at the postsynaptic membrane to ensure proper synaptic transmission.

## Introduction

It is well established that microtubule motor proteins are crucial for neuronal connectivity and function. In fact, mutations in cytoplasmic dynein, the dynein co-factor dynactin, or kinesin-1, result in a number of neurodegenerative disorders (Ebbing et al., 2008; Farrer et al., 2009; Inaki et al., 2010; Weedon et al., 2011; Tsurusaki et al., 2012; Brenner et al., 2018). The onset of these neurodegenerative disorders typically follows a loss in synaptic connectivity and function between the neuron and its target tissue, often another neuron or muscle cell. Studies on the function of dynein and kinesin motors in neuronal connectivity have focused primarily on their presynaptic role in axonal transport. Numerous studies have elucidated the role of microtubule motors in the anterograde transport of synaptic components from the cell body to the synaptic terminal, the retrograde transport of survival signals from the synapse to the cell body, and in the transport mechanisms that remove misfolded proteins and cellular debris from the axon (Okada et al., 1995; Bhattacharyya et al., 2002; Delcroix et al., 2003; Klopfenstein and Vale, 2004; Pack-Chung et al., 2007; Maday et al., 2012). Less well understood is the role of motor proteins on the postsynaptic side of neuronal connections.

The majority of excitatory glutamatergic synaptic transmission in the mammalian CNS occurs on dendritic spines of neurons. These specialized postsynaptic compartments protrude from the dendritic branches to synapse with other neurons. While the shafts of the dendrites are enriched in microtubules, dendritic spines are enriched in F-actin and were previously thought to be largely devoid of microtubules (Kaech et al., 1997; Kaech et al., 2001). However, more recent studies have demonstrated that microtubules do in fact enter the dendritic spines (Gu et al., 2008; Hu et al., 2008; Gu and Zheng, 2009; Jaworski et al., 2009). Studies examining microtubule motor function in dendrites have found that both dynein and kinesin-1 act to transport AMPA glutamate receptors into dendritic shafts (Kapitein et al., 2010; Heisler et al., 2014). In turn, myosin motors are thought to transport these receptors, and other cargoes, along the actin cytoskeleton to the membrane of dendritic spines (Wang et al., 2008; Esteves da Silva et al., 2015). The delivery of specific subsets of cargoes to dendritic spines may involve other kinesins. For example, the transport of synaptotagmin IV requires the kinesin-3 family motor KiF1A (McVicker et al., 2016), while multiple related kinesin-1s (e.g., KIF5A,B, and C) are differentially involved in the transport of RNA-binding proteins that contribute to spine morphogenesis, density, and plasticity (Zhao et al., 2020). It will be important to understand the full repertoire of microtubule-based motors in dendrites that contribute to synaptic function.

Drosophila Type 1 neuromuscular junction (NMJ) synapses are glutamatergic, containing ionotropic glutamate receptors that are functionally equivalent to AMPA receptors of the excitatory synapses in the mammalian CNS. In addition to receptors, the Drosophila structural, cytoskeletal, and membrane-associated components that organize and regulate the postsynaptic NMJ are functionally conserved with orthologous components in the postsynaptic dendritic spines of mammals. These include cell adhesion molecules that link the presynaptic and postsynaptic compartments (e.g., Neuroligins and NCAMs/Fas2), components of the postsynaptic density (e.g., PSD-95/Dlg), and regulators of the actin cytoskeleton (e.g. Adducin/Hts, and α/β-Spectrins)(Guan et al., 1996; Matsuoka et al., 1998; Pielage et al., 2006; Kohsaka et al., 2007; Banovic et al., 2010; Sun et al., 2011; Wang et al., 2011). Moreover, similar to mammalian dendritic spines, the postsynaptic side of the Drosophila NMJ is enriched in F-actin and thought to be largely devoid of microtubules, although only a subset of microtubules has previously been examined (Coyle et al., 2004; Ruiz-Canada et al., 2004; Ramachandran et al., 2009). Given the similarities in structure and function, the Drosophila NMJ is an excellent model to further probe the postsynaptic function of microtubule motor proteins.

Here, we investigated the function of microtubule motor proteins on the postsynaptic side of excitatory glutamatergic synaptic terminals in Drosophila. We report that cytoplasmic dynein accumulates on the postsynaptic side of NMJs. Our subsequent studies into postsynaptic dynein function suggest that dynein-based transport of a PI4P5 kinase, Skittles, to the postsynaptic NMJ regulates the organization of the postsynaptic spectrin-actin cytoskeleton. Additionally, postsynaptic dynein is required for synaptic growth. We further revealed that dynein, independent of Skittles, is required to organize glutamate receptor clusters at the postsynaptic NMJ, impacting synaptic transmission.

## Results

### Dynein localizes to the postsynaptic neuromuscular junction through its motor activity

In the skeletal muscle of third instar Drosophila larvae, we find that dynein has a unique, postsynaptic localization at neuromuscular junctions (NMJs) which has not been previously described. To investigate the postsynaptic function of motor proteins at the NMJ, we characterized the organization of the postsynaptic microtubule cytoskeleton and the distribution of the motor proteins cytoplasmic dynein and kinesin-1 within the postsynaptic muscle. We find that acetylated microtubules overlap with a postsynaptic density protein, β-Spectrin, at the NMJ (Figure 1A-A”), advocating for a potential postsynaptic role for microtubule motor proteins. Further, the heavy chain subunit of cytoplasmic dynein (Dhc64c) localizes to punctae in the postsynaptic muscle surrounding the presynaptic neuronal tissue at NMJs (Fig. 1B-B’’). Intriguingly, although dynein expression in larval skeletal muscles is ubiquitous, the localized accumulation of dynein punctae is specific to glutamatergic Type I synaptic terminals as we do not observe dynein accumulation at Type III peptidergic terminals (Figure 1C-C’), nor at Type II octopaminergic terminals. Similarly, we find that another subunit of the complex, dynein light intermediate chain (dlic), localizes to punctae surrounding the neuromuscular junction (Figure S1A-A’). Moreover, a punctate postsynaptic localization has been noted for Glued, the p150 subunit of the dynactin complex, an accessory complex to the dynein motor (Eaton et al., 2002). To confirm that the punctae we observe are specific to cytoplasmic dynein, we depleted the dynein heavy chain subunit by RNAi using the driver *M12-Gal4*, which expresses specifically in muscle 12 (Inaki et al., 2010). We then compared the staining of dynein in muscle 12 to the adjacent muscle 13. While Dhc64c puncta are observed in muscle 13, in muscle 12 Dhc64c is only present on the neuronal side of the NMJ (Fig. 1D-D’). Together, these results demonstrate that the punctate staining that we observe is dynein and muscle specific.

**Figure 1.**
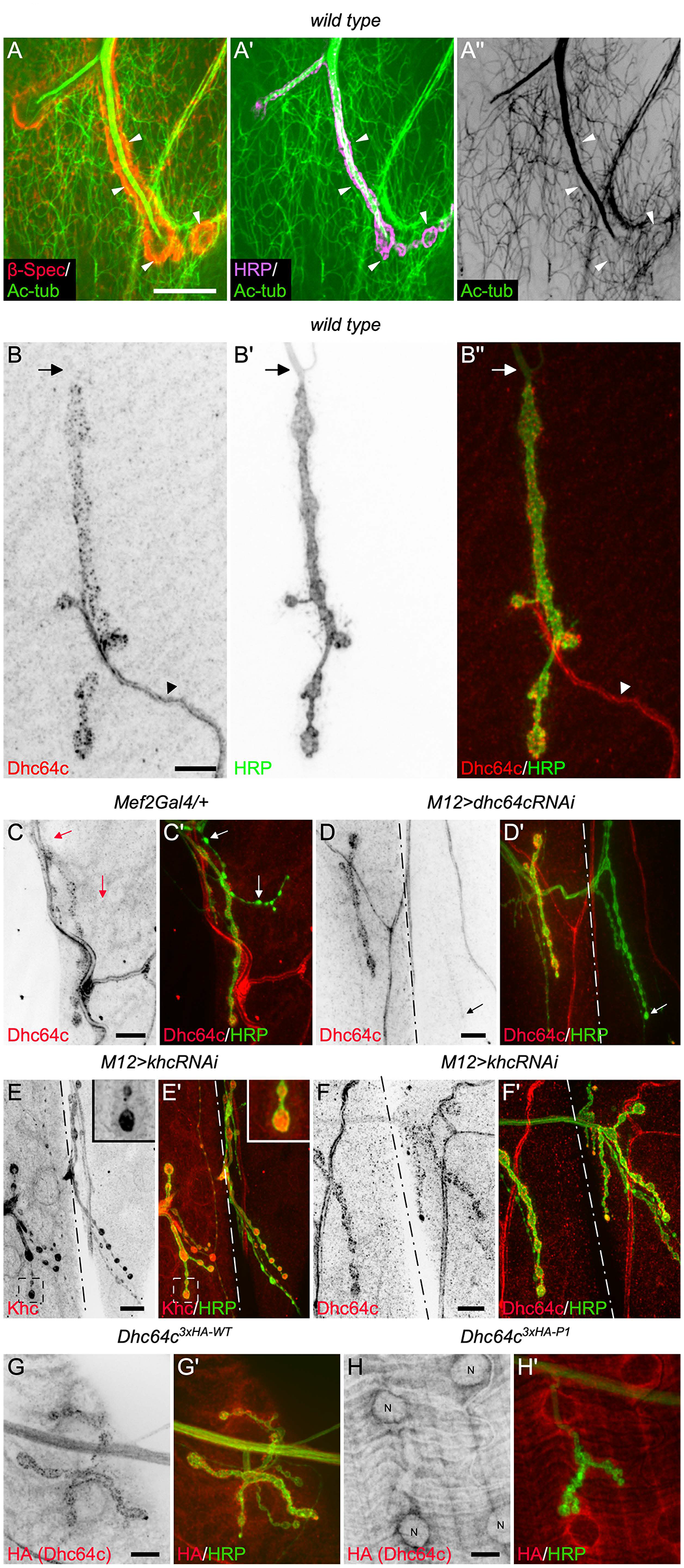
Cytoplasmic dynein has a punctate postsynaptic localization that requires its motor function. The microtubule cytoskeleton at the NMJ was visualized using an acetylated-α-tubulin antibody. β-spectrin localization marks the postsynaptic side of the NMJ **(A)**, while HRP labels neuronal membranes **(A’)**. A subset of microtubules overlap with postsynaptic β-spectrin, some of which are indicated by the white arrowheads **(A-A’’)**. The localizations of cytoplasmic dynein and kinesin-1 at the NMJ were assessed using antibodies that specifically recognize the dynein heavy chain subunit, Dhc64c, and the kinesin heavy chain subunit, Khc. **B-B’’)** Dynein heavy chain localizes to postsynaptic punctae that are in close apposition to the neuronal membrane labeled with HRP. The arrows indicate the position of the axon which lacks dynein punctae. The arrowheads indicate a tracheal branch in the muscle that also is enriched with dynein. **C-C’)** Dynein specifically localizes to glutamatergic Type I synaptic terminals. The arrows indicate a Type III peptidergic terminal that is lacking dynein punctae. **D-D’)** Depletion of dynein in muscle 12 (right side of the dotted line), demonstrates that these punctae are postsynaptic, and dynein specific. The left side of the line is muscle 13 for comparison. The arrow indicates visible presynaptic dynein staining when dynein is depleted from the muscle. **E-E’)** Kinesin-1 was depleted from muscle 12 (right side of the dotted line). Depletion of kinesin-1 in muscle 12 when compared to the endogenous staining of kinesin-1 in muscle 13 (left side of the line) shows that kinesin is enriched on the presynaptic side of the NMJ without an obvious postsynaptic enrichment. The boxed area on muscle 13 is shown as an inset, to show kinesin staining at the NMJ. **F-F’)** Depletion of kinesin-1 from muscle 12 (right side of the dotted line) does not affect the subcellular postsynaptic localization of dynein. **G-G’)** A 3xHA-tagged heavy chain subunit expressed under its endogenous promoter has a similar postsynaptic punctate distribution as observed with the Dhc64c antibody. **H-H’)** A 3xHA-tagged heavy chain subunit containing mutations in P-loop 1 (P1), required for ATP hydrolysis, does not localize to postsynaptic punctae, demonstrating the requirement of motor activity for dynein’s postsynaptic NMJ localization. Instead, dynein accumulation is visible surrounding the muscle nuclei, labelled with the letter N **(H)**. Scale, 10μm.

The plus-end directed motor protein, kinesin-1, often functions cooperatively with cytoplasmic dynein. In particular, within neurons dynein and kinesin-1 act in concert to regulate the transport of numerous cellular cargoes, including motor complexes themselves, up and down the axon (Martin et al., 1999; Pilling et al., 2006; Lloyd et al., 2012; Twelvetrees et al., 2016; Neisch et al., 2017). Additionally, kinesin-1 is involved in transport of glutamatergic receptor subunits in other systems (Hoerndli et al., 2013; Heisler et al., 2014), suggesting a potential postsynaptic function. To test this possibility, we examined the subcellular localization of kinesin-1 at NMJ using both a kinesin-1 specific antibody and a ubiquitously expressed kinesin-1-GFP protein. In contrast to the localization of dynein at the NMJ, we do not observe an accumulation at the postsynaptic NMJ of kinesin-1 (Figure 1E-E’, S1B-B’). Instead, kinesin-1 is enriched within the presynaptic terminals. As dynein and kinesin are known to functionally interact, we also asked if kinesin-1 was required for the postsynaptic NMJ localization of dynein. We depleted kinesin heavy chain (Khc) from muscle 12 and examined the effects of this depletion on dynein localization. We found that depletion of kinesin-1 does not impact the postsynaptic NMJ accumulation of dynein (Figure 1F-F’, S1D-D’), suggesting there is no interdependence of the motor proteins for dynein’s postsynaptic localization. Interestingly, we did observe halos of dynein staining around nuclei when kinesin is depleted from the muscle (Figure S1 compare C-C’ to D-D’), suggesting that kinesin and dynein may work cooperatively in other functions within the muscle. In fact, during embryogenesis both kinesin and dynein are reportedly required for nuclear cell positioning within the muscle and show interdependence in this process (Folker et al., 2014). Nonetheless, our results suggest that dynein has a unique postsynaptic localization and function that is independent of kinesin-1 activity.

We speculated that dynein’s localization at the NMJ was due to its motor activity. To test this prediction, we examined the localization of a previously characterized Dhc64c mutant protein (Dhc64c^P1-3HA^), defective in ATP hydrolysis and microtubule-based motility (Silvanovich et al., 2003). By comparison to a wildtype, epitope-tagged Dhc64c (Dhc64c^WT-3HA^), the P-Loop1 dynein mutant protein does not localize to postsynaptic punctae at the NMJ, demonstrating that dynein motor activity is required for its localization at the NMJ (Fig. 1G-H). This result also implies that at least a subset of the microtubules within the muscle are oriented with their minus-ends toward the NMJ, as dynein is a minus-end directed motor. We hypothesize that dynein’s motor activity is important for the transport and localization of postsynaptic proteins and/or RNAs required for NMJ functionality.

### Postsynaptic dynein is necessary for synaptic growth

Drosophila undergo a rapid growth period, increasing their body mass ∼200 fold over 4 days (Lilly and Duronio, 2005; Tennessen and Thummel, 2011). During this period of rapid growth, body wall muscles also grow significantly, with muscle surface area increasing ∼100 fold. In order to maintain synaptic homeostasis, the presynaptic terminal must grow as well, continuously adding synaptic boutons (reviewed in Menon et al., 2013). The disruption of several cytoskeletal components has been found to impede the homeostatic balance between presynaptic terminal and muscle growth, resulting in both decreased and increased growth of the presynaptic terminal as measured by the number of boutons in third instar larvae (Roos et al., 2000; Sherwood et al., 2004; Pielage et al., 2006; Pawson et al., 2008; Graf et al., 2011; Zhao et al., 2013; Blunk et al., 2014; Mao et al., 2014; Migh et al., 2018; Chou et al., 2020). Many of these effects have been attributed to cytoskeletal proteins acting within the neuron, while our knowledge of postsynaptic cytoskeletal elements that contribute to synaptic growth are limited (Pielage et al., 2006; Blunk et al., 2014).

Given the unique pattern of postsynaptic dynein accumulation at the NMJ, we asked if postsynaptic dynein is important for synaptic growth and NMJ morphology. We depleted dynein from the muscle by driving the expression of an RNAi line that targets the dynein heavy chain subunit, using the somatic muscle driver *Mef2-Gal4*. *Mef2-Gal4* is expressed in all somatic muscles from embryonic stage 7 onward (Ranganayakulu et al., 1996). To analyze the NMJs, we used the presynaptic membrane marker, horseradish peroxidase (HRP), and Discs Large (Dlg) which localizes postsynaptically. We quantified the number of synaptic boutons in third instar larvae, focusing on type 1b boutons at NMJ4 of abdominal segment A3 and at NMJ6/7 of abdominal segment A2, both well characterized NMJs that are commonly used for analysis of synaptic growth and function (Shen and Ganetzky, 2009; Xing et al., 2014; Han et al., 2020). To determine if there was a change in synaptic growth, we quantified the number of boutons per muscle area. Notably, we find that depletion of dynein within the muscle results in a significant reduction in the number of boutons, while having no significant impact on the muscle area (Figures 2, S2A-A’). These results indicate that postsynaptic dynein does indeed impact synaptic growth. One possibility is that dynein delivers and/or localizes one or several components postsynaptically at the NMJ to regulate synaptic terminal growth.

**Figure 2.**
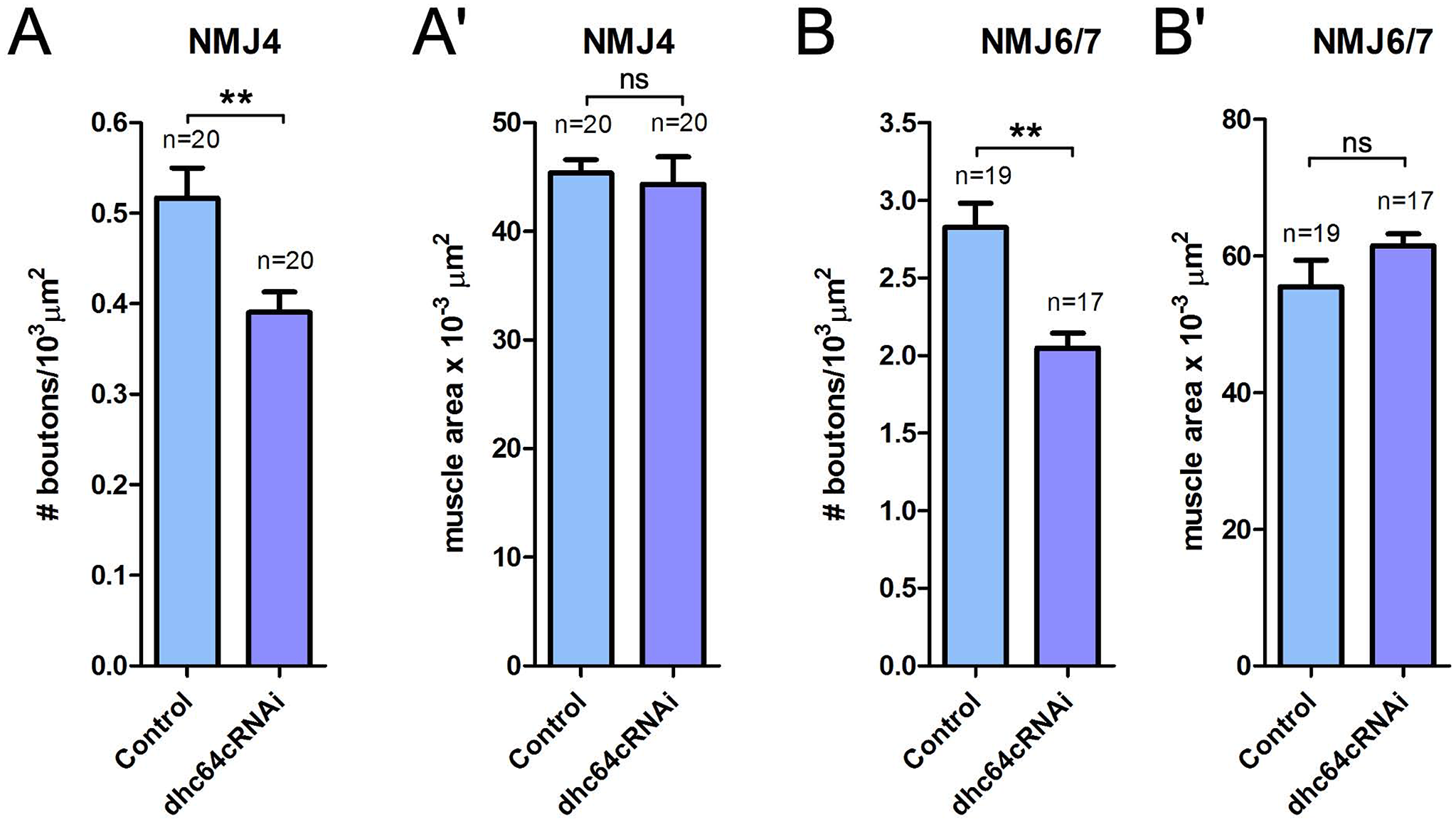
Postsynaptic dynein modulates synaptic growth. To probe if postsynaptic dynein influences synaptic growth, the number of boutons and the total muscle area were measured for muscle 4 and muscles 6/7. **A)** The number of boutons per muscle area at NMJ4 is significantly reduced when dynein is depleted. However, the muscle area is not significantly affected **(A’)**. **B)** Similarly, the number of boutons per muscle area at NMJ6/7 is significantly reduced when dynein is depleted. Again, the muscle area is not significantly affected by depletion of dynein in the muscle **(B’)**. n=number of NMJs analyzed. Unpaired two-tailed t-test results: **, P<0.005, ns=not significant.

### Postsynaptic dynein is required for proper glutamate receptor clustering

We have established that dynein’s localization is specific to glutamatergic synaptic terminals (Figure 1C-C’). We noted that the punctate distribution of dynein looked similar to that reported for glutamate receptors, and for this reason looked at the distribution of the two in relationship to one another. We find that the distribution of dynein aligns with the pattern of glutamate receptor subunits (Figure 3A-A’’), with some punctae showing an overlapping localization (yellow in inset 3A’’). We therefore asked if dynein is required for the localization of the glutamate receptor subunits at the NMJ. We postulated that dynein acts to transport subsets of receptors and/or receptor adaptors to the postsynaptic terminal, or alternatively may act to remove receptors from the terminal.

**Figure 3.**
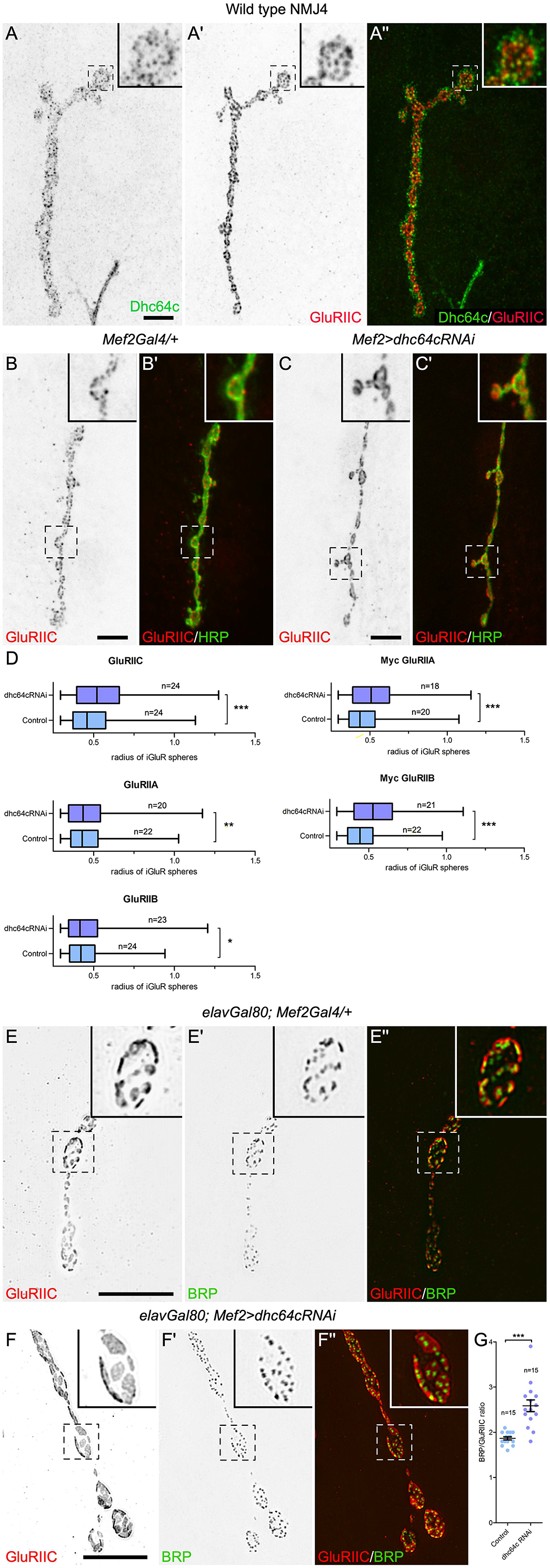
Dynein is required for ionotropic glutamate receptor clustering. **A-A”)** The punctate distribution of dynein, while somewhat broader, is similar to that of the glutamate receptor subunit GluRIIC at the NMJ. A single optical section through the NMJ is shown. In the inset, many GluRIIC positive punctae overlap with dynein punctae and appear yellow **(A’’)**. An example of the GluRIIC localization in a control animal **(B-B’)** and in a dynein depleted animal **(C-C’)** used for the analyses in **(D)**. HRP labels the presynaptic side of the NMJ, and single optical sections through the NMJ are shown. **D)** Boxplots of the radius of iGluR spheres in control animals, and those depleted of dynein by RNAi. n=number of NMJ analyzed. Unpaired, one-tailed t-test results: ***, P<0.0001, **, P<0.005, *, P<0.05. **E-E’’)** Structured illumination microscopy images of the GluRIIC subunit and Bruchpilot (BRP), a presynaptic active zone component, in control animals and in animals depleted of dynein within the muscle **(F-F’’)**. **G)** Quantification of the number of BRP, active zone punctae apposed to a postsynaptic glutamate receptor cluster for control animals and those depleted of muscle dynein. n=number of NMJ analyzed, each point on the graph is an individual NMJ. Unpaired, two-tailed t-test results: ***, P<0.0001. Boxed areas are shown as insets. Scale, 10μm.

In Drosophila, there are five glutamate receptor subunits. The receptor complexes are heteromeric tetramers comprised of three essential subunits; GluRIIC, GluRIID, and GluRIIE, that assemble with either GluRIIA, referred to as Type A receptors, or GluRIIB, referred to as Type B receptors (Qin et al., 2005). Changes in the quantity of Type A or Type B receptors result in physiological differences. For instance, an increase in Type A receptors elevates the postsynaptic response to the quantal release of neurotransmitter upon spontaneous fusion of a single synaptic vesicle (quantal size), while increasing levels of Type B receptors has the opposite affect (DiAntonio et al., 1999). Staufen, an RNA binding protein, is required for GluRIIA/GluRIIB levels at NMJ in Drosophila (Gardiol and St Johnston, 2014), and Staufen, in other systems, complexes with the dynein motor (Gagnon et al., 2013; Gershoni-Emek et al., 2016) suggesting dynein may also impact GluRIIA/GluRIIB levels through trafficking Staufen. We therefore examined the effects of dynein muscle depletion on both the quantity and localization of three of the receptor subunits, GluRIIC, GluRIIA, and GluRIIB. Surprisingly, we find that loss of dynein does not significantly impact the quantity of any of the receptor subunits at the NMJ relative to controls (Figure S2B). However, when we examine the localization of the glutamate receptor subunits, we observe that loss of dynein results in larger receptor clusters (Figure 3B-C, GluRIIC, Figs S2C-F, GluRIIA and GluRIIB). We quantified the receptor cluster size and found that all three subunits in the complex show a significant increase in size when dynein is depleted (Figure 3D). Because of the weak staining and higher background of the GluRIIA and GluRIIB antibodies, we also measured the receptor cluster sizes of Myc-tagged GluRIIA and GluRIIB expressed with the muscle specific promoter, Myosin heavy chain (Mhc). Again, we find a significant increase in cluster size when dynein is depleted compared to controls (Figures 3D, S2G-J). As dynein depletion significantly affects the size of receptor clusters for all three receptor subunits examined, these results suggest that dynein is required for the proper assembly/organization of the receptor fields at the NMJ.

To acquire higher resolution images of the glutamate receptor fields and the presynaptic active zones that are apposed to the receptor clusters, we used super resolution Structural Illumination Microscopy (SIM). We examined receptor clusters in control animals and those depleted of postsynaptic dynein (Figures 3E-F’’). Because we were examining how loss of postsynaptic dynein impacts both the presynaptic and postsynaptic tissue, we wanted to ensure that we were only disrupting the function of dynein within the muscle. To do this, we depleted dynein from the muscle using the *Mef2-Gal4* driver and co-expressed *elav-Gal80* to suppress any possible Gal4 expression within neurons. Glutamate receptor clusters were visualized using the GluRIIC receptor subunit, and active zones were visualize using Bruchpilot (BRP), an active zone scaffolding component required for active zone assembly and structural integrity (Kittel et al., 2006; Wagh et al., 2006). The receptor clusters are much larger in diameter when dynein is depleted from the postsynaptic muscle by comparison to control animals. Other investigators have reported ∼1 to 1.5 BRP punctae per glutamate receptor cluster in control animals (Pielage et al., 2006; Viquez et al., 2009; Wairkar et al., 2009; Hong et al., 2020). In our own analysis of SIM images, we find that there are on average 1.87±0.04 BRP punctae per glutamate receptor cluster in control animals. Significantly, this number was increased to 2.59±0.13 BRP punctae per glutamate receptor cluster when dynein is depleted from the muscle (Figure 3G). Taken together, these results suggest that organization of the glutamate receptor fields depends on dynein. The loss of postsynaptic dynein increases the receptor field size and impacts the presynaptic apposition of active zones.

### Postsynaptic dynein functions in modulating spontaneous synaptic transmission

To determine if there are physiological consequences resulting from postsynaptic dynein loss and the associated changes in glutamate receptor and active zone organization, we measured the spontaneous release of glutamate (mEJPs). Again, we depleted dynein postsynaptically in the muscle using the *Mef2-Gal4* driver and co-expressed *elav-Gal80* to suppress any possible Gal4 expression within neurons. Compared to controls, we find that depletion of dynein by RNAi results in both a significant increase in the frequency of mEJPs and increase in the amplitude of mEJPs (Figure 4A-B, D). In control animals, there are very few spontaneous release events with large amplitudes, with only 25.1% of events having amplitudes larger than 0.8mV. However, dynein depleted animals exhibit a greater number of events and increased amplitudes overall, with 40% of events having an amplitude larger than 0.8mV (Figure 4C). Increased amplitudes of mEJPs are observed for changes in postsynaptic glutamate receptor organization (Vautrin and Barker, 2003; Kittel et al., 2006; Pielage et al., 2006), and correlate with the changes we find in receptor clustering when dynein is depleted postsynaptically. Changes in the frequency of mEJPs is generally thought to be associated with presynaptic changes, as each peak is the result of neurotransmitter release upon fusion of a synaptic vesicle with the plasma membrane (reviewed in Bykhovskaia and Vasin, 2017). This result suggests that postsynaptic dynein mediates a transsynaptic effect at the NMJ.

**Figure 4.**
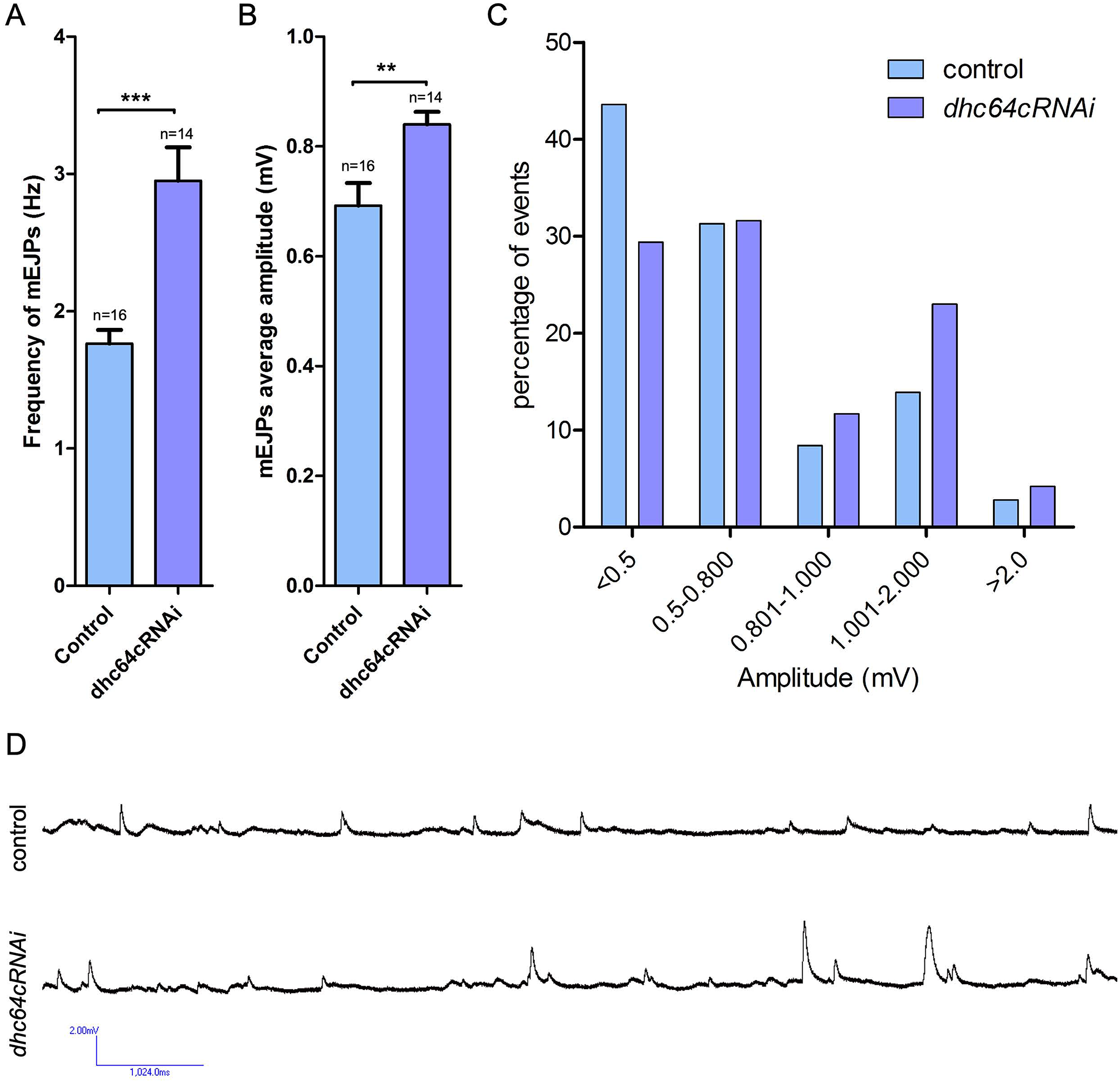
Postsynaptic dynein is important for controlling spontaneous synaptic transmission. Electrophysiology recordings were carried out to measure spontaneous release of glutamate containing vesicles (mEJPs) at NMJs in control animals and those depleted of dynein within the muscle. Muscle 6 of abdominal segment A2 and A3 were used for recordings. **A)** The frequency of mEJPs are significantly increased when dynein is depleted from the muscle. **B)** The average amplitude is also significantly increased when dynein is depleted from the muscle. **C)** A graph showing the percent distribution of events at increasing amplitudes in control animals and dynein depleted animals. **D)** Representative mEJP recordings from a control animal and a dynein muscle depleted animal. n=the number of recordings analyzed. Unpaired, two-tailed t-test results: ***, P<0.0001, **, P≤0.005.

### Dynein is necessary for the localization of membrane-associated postsynaptic proteins

To better understand how dynein functions to impact NMJ organization and physiology, we further analyzed the localization of two postsynaptic proteins, Dlg and β-Spectrin. Both Dlg and β-Spectrin are specific to glutamatergic synaptic terminals, localizing to the subsynaptic reticulum (SSR), an elaboration of the postsynaptic muscle cell membrane (Guan et al., 1996; Featherstone et al., 2001; Pielage et al., 2006). Dlg is a membrane-associated guanylate kinase (MAGUK) which is necessary for the formation of the SSR and scaffolding synaptic proteins at the SSR (Guan et al., 1996). Similarly, β-Spectrin is a part of the spectrin-actin network required for the organization of the SSR and postsynaptic densities (Pielage et al., 2006). To examine β-Spectrin localization, we developed a β-Spectrin specific antibody (Figure S3, see materials and methods). In wild type animals postsynaptic Dlg is closely juxtaposed to presynaptic boutons while β-Spectrin extends beyond the Dlg domain as visualized by immunolocalization studies (Figure 5A-A’’). Relative to Dlg and β-Spectrin, we find that dynein punctae lie just inside of the β-Spectrin localization domain (Figure 5B-B’’) and overlap with Dlg (Figure 5C-C’’). Significantly, both Dlg and the spectrin cytoskeleton have been implicated in influencing the size of the glutamate receptor field clustering (Pielage et al., 2006; Astorga et al., 2016), suggesting that dynein may be involved in the localization of one of these proteins.

**Figure 5.**
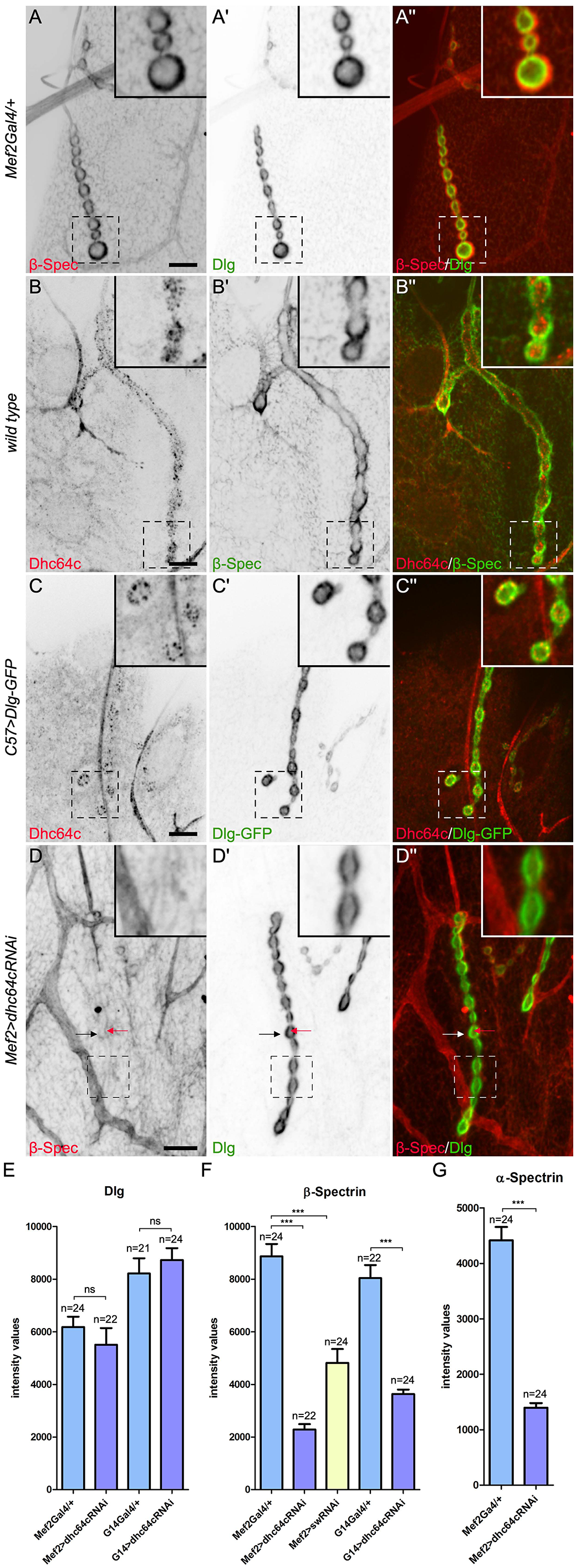
Dynein is required to organize the postsynaptic spectrin-cytoskeleton. The localization of dynein was examined with respect to two postsynaptic proteins, Dlg and β-Spectrin. **A-A”)** In control animals, Dlg and β-Spectrin partially overlap at the postsynaptic NMJ, with β-Spectrin localization extending beyond the localization of Dlg. **B-B’’)** Dynein punctae are found inside of the β-Spectrin localization domain at the NMJ. **C-C’’)** The localization of the dynein punctae overlaps with Dlg at the NMJ. **D-D’’)** Dynein depletion results in a disruption in the postsynaptic localization of β-Spectrin however it does not affect Dlg localization. The black arrows in (D-D”) indicate the postsynaptic side of the NMJ, where a low level of β-Spectrin staining is visible. The red arrows indicate the presynaptic β-Spectrin localization, which is not visible typically because of the greater intensity of postsynaptic β-Spectrin. **E)** Quantification of the intensity of Dlg at the NMJ in controls and when Dhc64c was depleted from the muscle, using two different muscle drivers. **F)** Quantification of the intensity of β-Spectrin at the NMJ in controls and when Dhc64c or the dynein intermediate chain (short wing, sw) were depleted from the muscle. **G)** Quantification of the intensity of α-Spectrin at the NMJ in controls and when Dhc64c was depleted from the muscle using the *Mef2-Gal4* driver. Boxed areas are shown as insets. Unpaired, two-tailed t-test results: ***, P<0.0001, ns=not statically significant. n=number of NMJs analyzed. Boxed areas are shown as insets. Scale, 10μm.

To address if dynein is required for the localization of postsynaptic Dlg or β-Spectrin, we depleted dynein by RNAi using the muscle drivers *Mef2-Gal4* and *G14-Gal4*. *G14-Gal4*, like *Mef2-Gal4,* is expressed in all somatic muscles, beginning at embryonic stages (Shishido 1998). Although dynein and Dlg have overlapping localizations, we find that when dynein is depleted from the muscle there is no change in the intensity of Dlg at the postsynaptic membrane (Figures 5D’, E). In contrast, the levels of β-Spectrin are significantly reduced when dynein is depleted (reduced by ∼55% with *G14-Gal4,* and by ∼74% with *Mef2-Gal4*; Figure 5D, F). Likewise, we find that the levels of α-Spectrin, which tetramerizes with β-Spectrin, are also significantly decreased (reduced by ∼68%; Figure 5G). Using an RNAi line against another dynein subunit, the dynein intermediate chain, known as short wing (sw) in Drosophila, we confirmed that depletion of dynein reduces postsynaptic β-Spectrin levels (reduced by 46%, Figure 5F), impacting the postsynaptic spectrin cytoskeleton at the NMJ.

Transmembrane cell adhesion molecules promote synapse assembly and postsynaptic organization, including the organization of the spectrin cytoskeleton. To determine if dynein organizes the spectrin cytoskeleton through localizing cell adhesion molecules at the NMJ, we examined two transmembrane cell adhesion proteins, Neuroligin 1 (Nlg1) and Teneurin-m (Ten-m). Similar to dynein, both Ten-m and Nlg1 are required for synaptic growth and organization of the spectrin cytoskeleton, but not postsynaptic Dlg localization (Banovic et al., 2010; Mosca et al., 2012; Mozer and Sandstrom, 2012; Xing et al., 2018). To quantify the changes in Nlg at the NMJ, we used two different Nlg1 specific antibodies as well as GFP-tagged Nlg1, which we expressed specifically in the muscle (Banovic et al., 2010; Mozer and Sandstrom, 2012). For assessing Ten-m levels at the NMJ, we used a specific Ten-m antibody. For both cell adhesion molecules, we observe no reduction in the levels of these proteins following depletion of postsynaptic dynein (Figures 6A-B, S4A-K’). Our results suggest that dynein does not function by altering the trafficking of the transmembrane cell adhesion molecules Ten-m or Nlg1.

**Figure 6.**
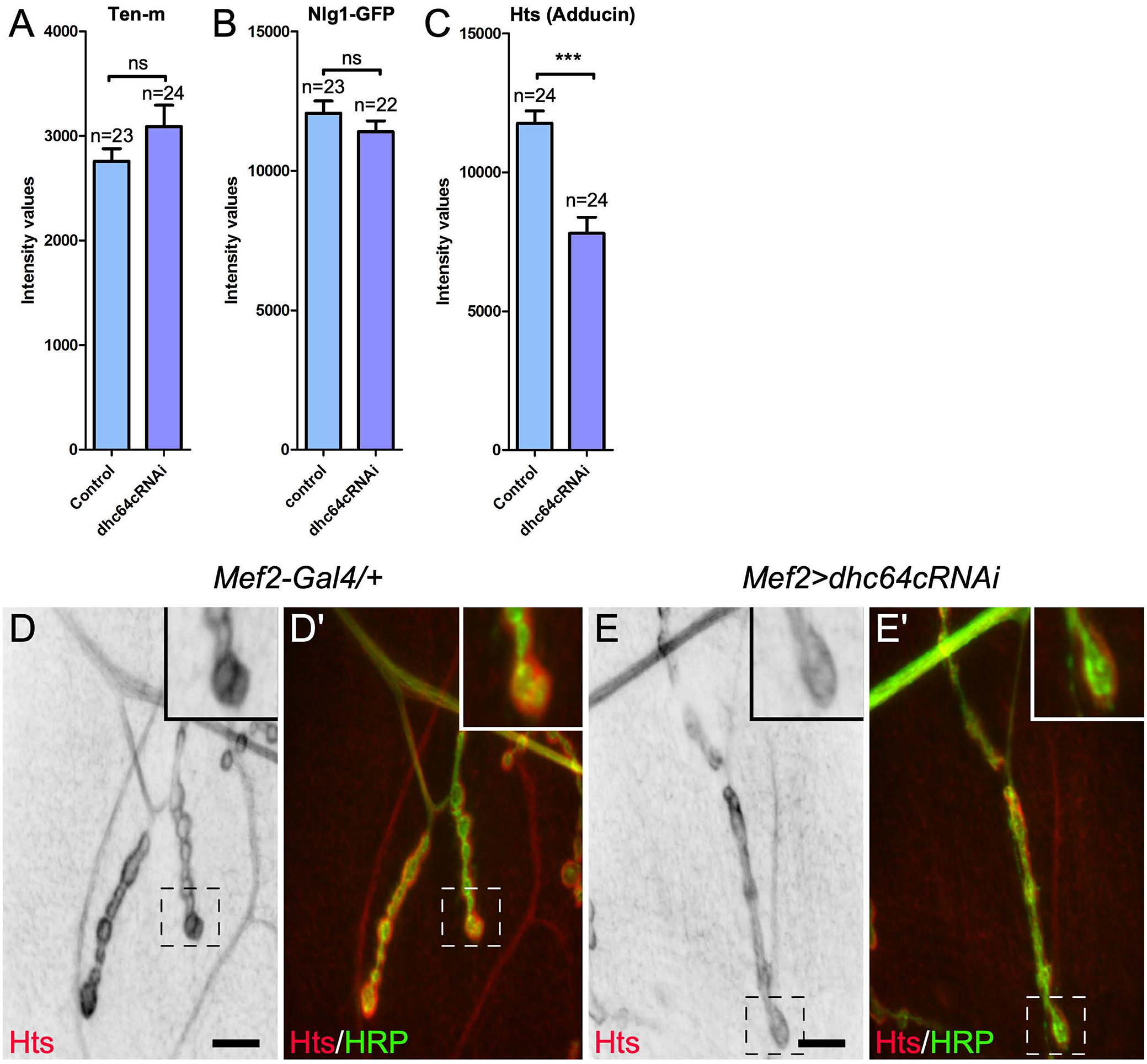
Dynein is required for the localization of Adducin at the NMJ, but not for the localization of cell-adhesion molecules Ten-m and Nlg1. To determine how dynein was impacting the spectrin-cytoskeleton, the localization of postsynaptic components that influence the spectrin-cytoskeleton were examined after dynein depletion from the muscle. No significant changes were observed for Ten-m or Nlg-GFP localization, while Adducin levels were reduced. Quantification of Ten-m **(A)**, Nlg-GFP **(B)**, and Hts (Adducin) **(C)** intensity levels at NMJ4. **D-D’)** The localization of Hts (Adducin) at the NMJ in a control animal. **E-E’)** The localization of Hts is diminished when dynein is depleted from the muscle. Unpaired, two-tailed t-test results: ***, P<0.0001, ns=not statically significant. n=number of NMJs analyzed. Boxed areas are shown as insets. In all images, HRP labels the presynaptic side of the NMJ. Scale, 10μm.

We further evaluated whether dynein impacts the recruitment of the spectrin associated protein Adducin. The membrane-associated protein Adducin organizes the spectrin cytoskeleton and is known as Hu-li tai shao (Hts) in Drosophila. Adducin is a cytoskeletal protein that links α/β-Spectrin tetramers to actin filaments (Gardner and Bennett, 1987; Mische et al., 1987). At Drosophila NMJs Adducin localizes to both the presynaptic and postsynaptic NMJ (Pielage et al., 2011; Wang et al., 2011). As Adducin links the spectrin cytoskeleton to actin at the NMJ, we asked if levels of Adducin are perturbed by the loss of dynein within the muscle. Indeed, we find that the levels of postsynaptic Adducin are significantly reduced when dynein is depleted from the muscle (reduced by ∼34%; Figure 6C-E’). Thus, dynein may impact the localization of the spectrin cytoskeleton by modifying the levels of Adducin at the NMJ, either through trafficking Adducin or regulating an upstream factor that is required for Adducin localization at the NMJ.

### Dynein is required for the localization of Sktl and PIP_2_ production at the NMJ

The loss of membrane-associated postsynaptic proteins at the NMJ led us to ask if the composition of the postsynaptic membrane is altered when dynein is depleted postsynaptically. The phospholipid membrane component, phosphatidylinositol 4,5-bisphosphate (PIP_2_) is known to accumulate postsynaptically at the NMJ (Wang et al., 2014). We therefore examined the levels of PIP_2_ at the postsynaptic membrane using a reporter that contains the pleckstrin homology domain from PLCdelta which binds PIP_2_ (Tan et al., 2014; Wang et al., 2014; Li et al., 2020). We find that depletion of dynein in the muscle significantly reduces the levels of PIP_2_ at the postsynaptic membrane (reduced by ∼38%, Figure 7A-C). The reduction in PIP_2_ levels at the NMJ is consistent with the reduction in Hts/Adducin and β-Spectrin that we observe at the NMJ (Figures 5D,F, 6C-E’). as both Adducin and β-Spectrin bind PIP_2_ (Gardner and Bennett, 1987; Mische et al., 1987; Das et al., 2008; Wang et al., 2014).

**Figure 7.**
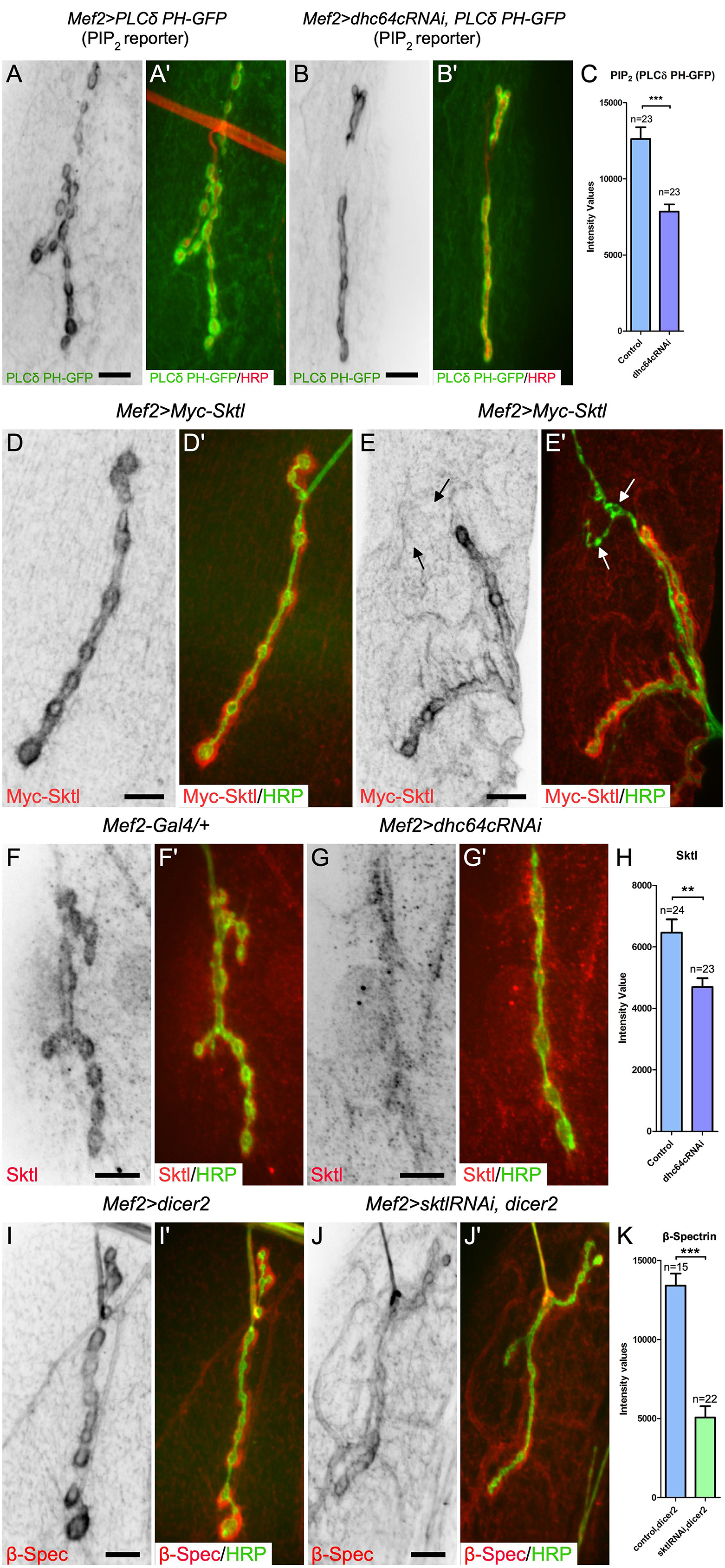
Dynein is essential for the localization of Skittles kinase to the NMJ for production of PI(4,5)P*_2_* locally at the postsynaptic NMJ membrane. To examine the distribution of PIP_2_ at the postsynaptic NMJ membrane the PIP_2_ reporter, PLC*δ*-PH-GFP, was expressed in the muscle. **A-A’)** PIP_2_ localization is enriched at the postsynaptic membrane. **B-B’)** Depletion of dynein in the muscle reduced the postsynaptic PIP_2_ membrane enrichment. **C)** Quantification of the postsynaptic membrane intensity of PIP_2_ (PLC*δ*-PH-GFP) in control animals and those depleted of dynein by RNAi. n=number of NMJ analyzed. **D-D’)** Localization of 6x-Myc-tagged Skittles (Sktl) expressed in the muscle is enriched at the postsynaptic NMJ. **E-E’)** Skittles localization is specific to glutamatergic Type I synaptic terminals. The arrows indicate a Type III peptidergic terminal that is lacking Skittles. **F-F’’)** A Skittles specific antibody shows a similar postsynaptic localization pattern at the NMJ. **G-G’’)** The postsynaptic localization of Skittles is lost when dynein is depleted from the muscle. **H)** Quantification of the postsynaptic membrane intensity of Skittles in control animals and those depleted of dynein by RNAi. Postsynaptic β-spectrin levels in dicer2 expressing control animals **(I)** and those depleted of Skittles **(J)**. **K)** Quantification of the postsynaptic membrane intensity of β-spectrin in control animals and those depleted of Skittles by RNAi, shows a statistically significant decrease in Skittles depleted animals. n=number of NMJ analyzed. Unpaired, two-tailed t-test results: ***, P<0.0001, ** P<0.005. In all images, HRP labels the presynaptic side of the NMJ. Scale, 10μm.

PIP_2_ is produced by the conversion of PI(4)P into PIP_2_ by Phosphatidylinositol 4-Phosphate 5-kinases, of which there are two in Drosophila, PIP5K59B and Skittles (Hassan et al., 1998; Chakrabarti et al., 2015). A subunit of the dynein complex, the dynein light intermediate chain (dlic), was recently found to form a complex with Skittles kinase (Jouette et al., 2019). Interestingly, we find that similar to dynein localization, Myc-tagged Skittles localizes to neuromuscular junctions and is also specific to glutamatergic Type I synaptic terminals (Figure 7D-E’). Similarly, a Skittles specific antibody localizes to the postsynaptic side of Type I terminals (Figure 7F), and this localization is disrupted by the loss of dynein (reduced by ∼27%; Figure 7G, H). Together these results suggest that dynein is required to transport Skittles to the NMJ and that Skittles localization is required to locally produce postsynaptic PIP_2_. These results further suggest that postsynaptic PIP_2_ is required for binding Adducin and/or β-Spectrin to organize the spectrin cytoskeleton. To substantiate this model, we asked if Skittles was required at the NMJ for β-Spectrin postsynaptic organization. Using expression of Skittles RNAi in the muscle, we first confirmed that a RNAi line depletes Skittles using the muscle 12 specific driver, *M12-Gal4* (Figure S5A-B’’). After confirming that the RNAi line was effective at knocking down Skittles, we then asked if depletion of Skittles in the muscle, using *Mef2-Gal4*, changed the levels of postsynaptic β-Spectrin. Consistent with a requirement of Skittles in organizating of the spectrin-cytoskeleton, we observe a significant decrease in the levels of postsynaptic β-Spectrin (reduced by ∼62%; Figure 7I-K). Together our data support a model in which dynein actively transports Skittles to the postsynaptic NMJ where it locally facilitates PIP_2_ production. In turn, PIP_2_ contributes to organizing the postsynaptic spectrin cytoskeleton.

### Dynein mediates tethering of glutamate receptor clusters independent of Skittles

Both PIP_2_ and phosphatidylinositol-(3,4,5)-triphosphate (PIP_3_) phospholipids have been implicated in the localization of AMPA receptors at the postsynaptic membrane (Arendt et al., 2010; Seebohm et al., 2014). As dynein is required for both the localization of Skittles at the NMJ and proper glutamate receptor clustering, we tested whether Skittles, and hence PIP_2_ membrane levels are required to organize glutamate receptor clusters. We depleted Skittles from the muscle using the *Mef2-Gal4* driver and examined the effects of loss of Skittles on GluRIIC localization at the NMJ. Unlike with the depletion of dynein, postsynaptic depletion of Skittles results in an overall increase in GluRIIC levels at the membrane (Figure 8C). Additionally, there is increased cytoplasmic GluRIIC signal throughout the muscle, with a noted decrease in extrasynaptic GluRIIC punctae (Figure 8B, S5C compared to 8A, S5D). One interpretation is that reducing Skittles/PIP_2_ in the muscle results in a decreased turnover/removal of GluRs from the synaptic terminal, and this may account for a decrease in extrasynaptic GluRIIC punctae. Previously, it has been shown that glutamate receptors are removed from established postsynaptic densities, but at a low rate (Rasse et al., 2005). The only Skittles RNAi line that we found to be effective in depleting Skittles from the muscle also targets the other PIP5K in Drosophila, PIP5K59B. It is formally possible that PIP5K59B knockdown also contributes to the increased GluRIIC membrane postsynaptic membrane levels and decreased GluRIIC extrasynaptic punctae that we observe. More work is needed to resolve whether the two kinases produce distinct pools of PIP_2_ (Kolay et al., 2016).

**Figure 8.**
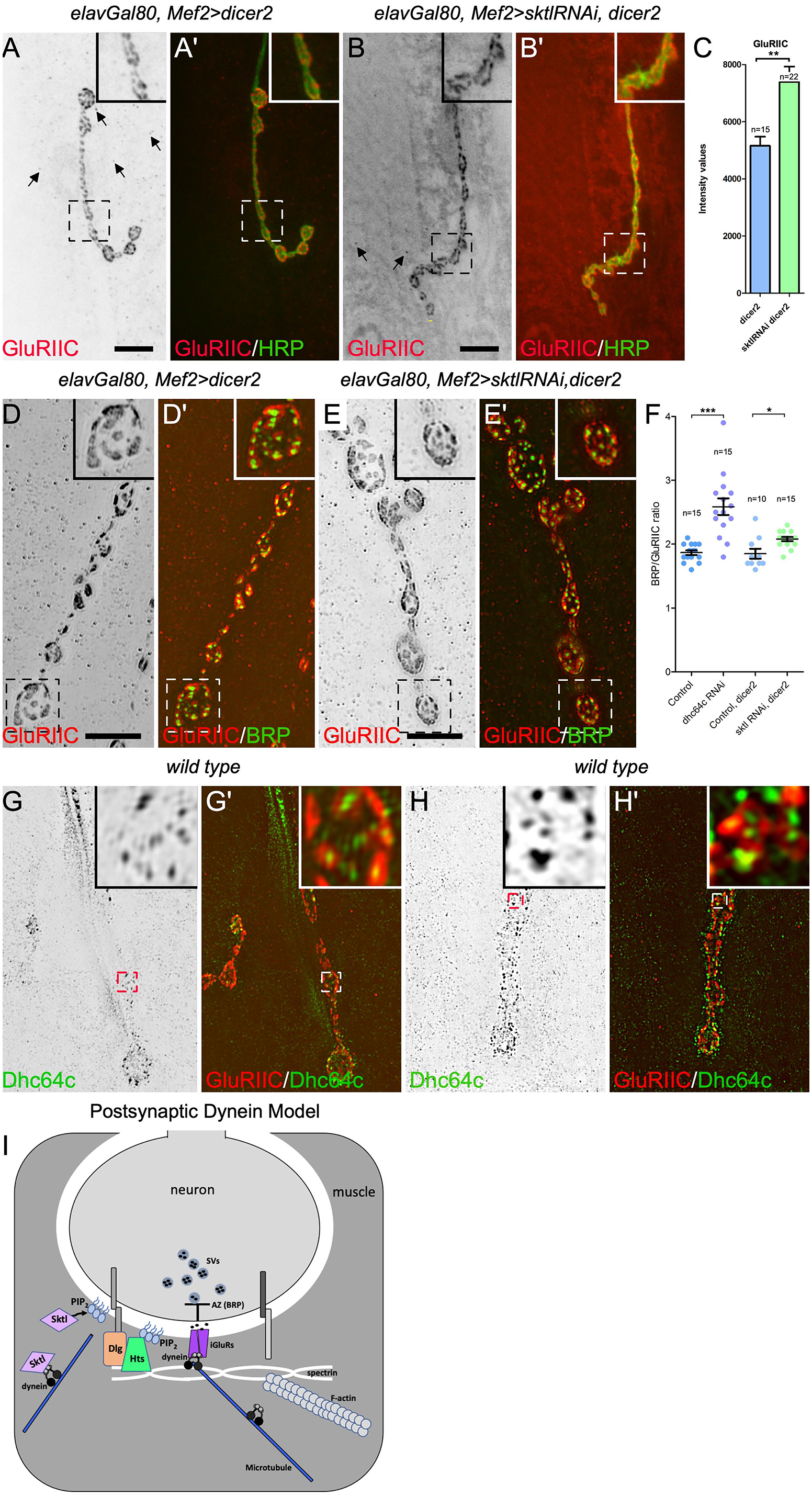
Skittles does not substantially contribute to glutamate receptor clustering while dynein punctae localize to the center of glutamate receptor clusters. Compared to controls **(A-A’)**, depletion of postsynaptic Sktl by RNAi results in increased cytoplasmic GluRIIC staining in the muscle and decreased extra synaptic GluRIIC punctae **(B-B’)**. Arrows indicated some of the extra synaptic punctae in **(A,B)**. **C)** Depletion of postsynaptic Sktl results in increased GluRIIC at the membrane compared to controls. Structured illumination microscopy was used to image and analyze GluRIIC clusters and their apposition to presynaptic BRP punctae. GluRIIC receptor cluster size, and their apposition to presynaptic Bruchpilot (BRP) punctae appear to be similar in controls **(D-D’)** and Sktl depleted animals **(E-E’)**. **F)** Quantification of the number of BRP, active zone, punctae apposed to a postsynaptic glutamate receptor cluster for control animals and those depleted of postsynaptic Sktl in comparison to the data shown in Figure 3H for dynein depletion, replicated again here for a side-by-side comparison. **G-H’)** Structured illumination microscopy was used to image glutamate receptor clusters and dynein together at NMJ to better determine the relationship between the two components. Two examples of wild type synaptic terminals are shown. GluRIIC labels glutamate receptor clusters, and Dhc64c labels the dynein heavy chain. In the merged image insets dynein often localizes to the center of the glutamate clusters **(G’,H’)**. **I)** A model for how dynein functions on the postsynaptic side of the NMJ to transport Sktl for local production of PIP_2_ and organization of the postsynaptic spectrin cytoskeleton, and stabilization/organization of glutamate receptor clusters. n=number of NMJ analyzed, each point on the graph is an individual NMJ. Unpaired, two-tailed t-test results: ***, P<0.0001, **, P<0.005, *, P<0.05. Boxed areas are shown as insets. Scale, 10μm.

To further characterize changes in the glutamate receptor fields, we compared high resolution SIM images of the GluRIIC receptor subunit and presynaptic active zones in controls, and following depletion of Skittles or dynein in the postsynaptic muscle. In contrast to the dynein depletion, the depletion of Skittles, does not result in enlarged glutamate receptor fields (Figures 3F-G compared to Figure 8D-E). In control animals, expressing only dicer2, we found that there are on average 1.85±0.07 BRP punctae per glutamate receptor cluster, similar to what we observed in our previous control samples (Figure 8F). When we depleted Skittles, we observed a slight increase from dicer2 controls to an average of 2.08±0.03 BRP punctae per receptor cluster. This difference is statistically significant, however not as great as the difference that we observe when dynein is depleted from the muscle (Figure 8F). This suggests that while dynein-mediated transport of Skittles and the postsynaptic accumulation of Skittles/PIP_2_ may contribute to glutamate receptor levels at the postsynaptic membrane, it cannot fully account for the receptor clustering defects we observe when dynein is depleted. Consistent with this interpretation, and in contrast to the postsynaptic loss of dynein (Figure 4), the loss of Skittles does not perturb synaptic transmission (Hassan et al., 1998). Together these results suggest that dynein is important to localize Skittles to the NMJ for the organization of the spectrin cytoskeleton, but that dynein functions largely independent of Skittles to organize the glutamate receptor clusters at the NMJ.

To better understand how dynein acts to organize glutamate receptor clusters at the NMJ, we analyzed the localization of the dynein heavy chain subunit, Dhc64c, and the glutamate receptor subunit, GluRIIC, using SIM. As we observed previously, the GluRIIC staining often forms donut-like structures. Remarkably, dynein localized to the center of many of the glutamate receptor clusters (Figure 8G-H’). The localization is very similar to that of BRP which localizes to the active zone, but on the presynaptic side of the junction at the center of the glutamate receptor clusters (Figures 3E’’ and 8D’). This pattern of localization together with the increased size of receptor clusters that results following postsynaptic depletion of dynein suggests that dynein may act to position the localization of glutamate receptors on the postsynaptic side of the NMJ.

## Discussion

Neuronal connectivity, maintained through pre- and post-synaptic connections, is crucial for cognitive and motor functions. Mutations within the motor protein complex, cytoplasmic dynein, and its accessory complex dynactin, are linked to a number of neurological disorders (Chevalier-Larsen et al., 2008; Lloyd et al., 2012; Moughamian and Holzbaur, 2012; Fiorillo et al., 2014; Hoang et al., 2017; Konno et al., 2017; Honda et al., 2018; Shen et al., 2018; Becker et al., 2020). While the role of microtubule motor proteins in neuronal transport and synaptic function is widely recognized, the postsynaptic roles of these proteins have been largely understudied. Here we uncover postsynaptic functions for cytoplasmic dynein in synaptic growth, the organization of glutamate receptor clusters, and the organization of postsynaptic spectrin cytoskeleton at the NMJ. Together our results suggest a model in which dynein has multiple separable functions at the postsynaptic NMJ. First, dynein acts to transport Skittles to the membrane, driving local PIP_2_ production necessary for the organization of the spectrin cytoskeleton. Second, dynein organizes glutamate receptor clusters at the membrane, potentially acting as a tether to stabilize their localization (Figure 8I).

### Postsynaptic Dynein vs Kinesin-1 at the NMJ

We discovered that postsynaptic dynein localization is specific to synaptic terminals that release the glutamate neurotransmitter. Conversely, we do not observe enrichment of the conventional kinesin, Kinesin-1, at the postsynaptic NMJ. While Kinesin-1 does seem to be required for the localization of a pool of dynein in the muscle, it does not affect dynein’s localization at the NMJ. In addition to Kinesin-1, there are 25 kinesin or kinesin-like proteins in Drosophila (Goshima and Vale, 2003; Hirokawa, 2012). Whether one of these other kinesins functions cooperatively with dynein at the postsynaptic NMJ remains to be determined.

### Dynein as a regulator of synaptic terminal growth

The rapid skeletal muscle growth that occurs during larval development is accompanied by coincident synaptic terminal growth to maintain stable physiological function. We find that dynein is required for growth of the synaptic terminal (Figure 2). Several components, both pre- and postsynaptic, display transsynaptic influences and can affect synaptic NMJ growth. These include cytoskeletal components, cell adhesion molecules, and signaling components (Marques et al., 2002; Packard et al., 2002; McCabe et al., 2003; Pielage et al., 2006; Li et al., 2007; Koch et al., 2008; Miech et al., 2008; Banovic et al., 2010; Mozer and Sandstrom, 2012). While morphological plasticity, such as NMJ terminal bouton number, and physiological plasticity, such as the regulation of neurotransmitter release, are thought to be linked, recent studies have revealed that this is not always the case (Kidd and Lieber, 2016; Borczyk et al., 2019; Goel et al., 2019). When dynein is depleted postsynaptically, we see both synaptic growth defects and physiological defects. It will be important in the future to determine whether homeostatic adaption mechanisms compensate for the synaptic terminal growth defects observed for postsynaptic dynein depletion.

The mechanism for dynein’s impact on synaptic terminal growth is yet to be determined. One possible mechanism is through dynein’s impact on PIP_2_ levels at the membrane. We have shown that depletion of dynein results in the reduction of PIP_2_ at the postsynaptic membrane (Figure 7). Among the many functions of PIP_2,_ its role in exocytosis of secretory vesicles is crucial in numerous physiological functions. For example, on the presynaptic side of neuronal connections the exocytosis of neurotransmitter containing vesicles is dependent on PIP_2_ (Grishanin et al., 2004; Bradberry et al., 2019). Additional work in MCF-7 cells has shown that PIP_2,_ and some of the isoforms of PIP5K, transiently accumulate specifically at sites of exocytosis (Stephens et al., 2020). In turn, accumulation of PIP_2,_ and PIP5K (i.e., Skittles), may function during postsynaptic exocytosis of Gbb, the BMP ligand. The BMP pathway is the major retrograde signaling pathway implicated in synaptic terminal growth at the NMJ (McCabe et al., 2003; Hoover et al., 2019). Loss of muscle derived Gbb results in undergrowth of the synaptic terminal (McCabe et al., 2003). The reduction of PIP_2_ levels following dynein depletion, may affect exocytosis of Gbb, or another unidentified molecule, from the muscle that is important for synaptic growth.

The impact of PIP_2_ on postsynaptic exocytosis may reflect its role as an important regulator of actin cytoskeleton dynamics. PIP_2_ interacts with multiple actin-binding domain proteins and promotes F-actin assembly, potentially driving the extension of the plasma membrane and exocytosis (reviewed in Katan and Cockcroft, 2020; Miklavc and Frick, 2020). Moreover, defects in the spectrin and actin cytoskeletons are common features of mutations exhibiting reduced synaptic terminal growth. For example, postsynaptic loss of each of the cell adhesion molecules Nlg1 and Ten-m result in both a synaptic terminal growth defect and a reduction in α/β-Spectrin (Mosca et al., 2012; Xing et al., 2018). In this study, we observe a correlation between reduced synaptic growth and a reduction in the levels of the F-actin binding proteins β-Spectrin, α-Spectrin, and Adducin following the depletion of postsynaptic dynein (Figures 5, 6). Consistent with these results, loss of postsynaptic β-Spectrin also results in decreased bouton numbers at the synaptic terminal (Pielage et al., 2006). Our results show that dynein is necessary for PIP_2_ accumulation at the NMJ, and we hypothesize that the differential affinities of distinct actin-binding proteins for PIP_2_ (Senju et al., 2017) in turn may regulate the underlying actin cytoskeleton and exocytosis.

### Skittles trafficking modulates PIP2 functions at the NMJ

Localization of the PIP4P5 kinase, Skittles, and its specific role at glutamatergic neuromuscular junctions has not previously been described. Our results show that Skittles localizes specifically to the postsynaptic side of glutamatergic synaptic terminals, and that this localization is dependent on dynein (Figure 7). Taken together, the localization of Skittles and accumulation of PIP_2_ (Wang et al., 2014), suggests that Skittles functions locally at the postsynaptic membrane to produce PIP_2_ at glutamatergic NMJs. PIP_2_, which primarily localizes to the plasma membrane, can be produced from either the precursor PI(4)P by PI4P5 kinases or the precursor PI(5)P by PI5P4 kinases. Interestingly, these kinases appear to produce distinct pools of PIP_2_ that carry out different cellular functions (Mayinger, 2012; Kolay et al., 2016). There is significant evidence for the actions of PIP_2_ at presynaptic sites, but knowledge of PIP_2_ function postsynaptically is limited. However, recent work suggests that PIP_2_ is a substrate for generation of inositol triphosphate (IP_3_) and diacylglycerol (DAG), key signaling elements underlying homeostatic plasticity in neurons (James et al., 2019). Thus dynein-mediated transport and localization of Skittles to the postsynaptic membrane of glutamatergic synapses may impact IP_3_/DAG signaling and affect transsynaptic signaling.

PIP_2_ also interacts with multiple membrane and cytosolic proteins to regulate a number of cellular processes in neurons. As discussed above, actin regulatory proteins may be differentially recruited to localized pools of PIP_2_ to control cytoskeletal polymerization and organization, which in turn can facilitate postsynaptic exocytosis involved in retrograde signaling. Furthermore, similar to the well characterized role for PIP_2_ in recycling of synaptic vesicles at presynaptic sites, the accumulation of PIP_2_ may also facilitate endocytosis at postsynaptic sites. Consistent with this interpretation, Skittles postsynaptic depletion results in an increase in overall GluRIIC accumulation at the membrane, and a coincident decrease in the number of extrasynaptic glutamate punctae within the muscle cell cytoplasm (Figure 8). Notably, increased intensity of GluRII receptor subunits at the NMJ is reported after postsynaptic inhibition of Shibire, the Drosophila dynamin GTPase known to mediate endocytosis (Nahm et al., 2010). This regulated trafficking and control of receptor abundance at the synaptic membrane is proposed to contribute to activity-induced changes in synaptic transmission (Malinow and Malenka, 2002; Bredt and Nicoll, 2003; Newpher and Ehlers, 2008). In this regard, dynein transport of Skittles may modulate local PIP_2_ levels and so regulate the endocytosis of receptor subunits.

### Cortical dynein as a tether for GluR clusters

At the NMJ, dynein localizes as punctae at the cortex of the postsynaptic cell membrane. These dynein punctae are closely associated with the center of glutamate receptor clusters that are arranged in a defined pattern, suggesting that the dynein motor complexes may act as a novel tether to organize glutamate receptors (Figure 8). Our results demonstrate that cortical dynein functions in the organization of the GluR receptor clusters at the postsynaptic membrane. Numerous cellular and developmental studies have implicated cortical dynein in the tethering of microtubule plus-ends and spindle orientation that underlies establishment of cell polarity, as well as the remodeling of cellular junctions during morphogenesis (Ligon and Holzbaur, 2007; Kotak et al., 2012; Lu and Prehoda, 2013; di Pietro et al., 2016; Rodriguez-Garcia et al., 2018). For example, in the case of epithelial adherens junctions, dynein localized at sites of cell-cell contact pulls on microtubules and the cytoskeleton to organize the interface between cells and to polarize the transport of cellular factors required at the specialized membrane junctions (Ligon et al., 2001; Ligon and Holzbaur, 2007). Similar to the adherens junctions, the immunological synapse (IS) is another site of cell-cell contact where dynein functions as a cortical tether (Combs et al., 2006; Martin-Cofreces et al., 2008; Hashimoto-Tane et al., 2011; Martin-Cofreces and Sanchez-Madrid, 2018). The IS exhibits several parallels to the Drosophila NMJ. Both at the IS and the NMJ dynein functions to organize the proper clustering of receptors in the membrane, and this organization is critical for proper receptor signaling (this work; Hashimoto-Tane et al., 2011). At the IS, PIP_2_ is cleaved to generate IP_3_ and DAG. Strikingly, DAG subsequently binds to and recruits dynein to the T cell membrane of the IS (Quann et al., 2009; Liu et al., 2013). Intriguingly, at the NMJ, PIP_2_ is also cleaved and is reported to produce the canonical second messengers DAG and IP_3_ that are integral to PHP (James et al., 2019). We suggest that similar to the IS, DAG produced from PIP_2_ at the postsynaptic NMJ may recruit dynein to the membrane.

While we report for the first time a postsynaptic role for dynein in Drosophila NMJs, dynein has previously been known to bind to mammalian NCAM180, a homophilic neural cell adhesion molecule localized on both sides of neurological synapses. Dynein interacts with NCAM180 and this interaction increases cell-cell adhesion and tethering of plus-end microtubules (Perlson et al., 2013). Moreover, when the NCAM180/dynein interaction is disrupted the density of active synapses is decreased, but the individual synapses were also much broader (Perlson et al., 2013). This broadening of the synapses may be similar to what we observe as an enlargement of the glutamate receptor clusters, but on the postsynaptic side, at the drosophila NMJ. Thus, NCAM180 is an alternative partner for recruiting dynein to the synapse. The Drosophila Fas2 isoforms are orthologous to the NCAMs present at the mammalian neurological synapse (Beck et al., 2012; Neuert et al., 2020) and may similarly direct the cortical recruitment of dynein that in turn interacts with and organizes receptor clusters of the postsynaptic membrane.

Our work has revealed novel postsynaptic dynein functions in synaptic terminal growth, glutamate receptor organization and the regulation of the postsynaptic cytoskeleton. To elucidate how dynein acts in these novel functions will require live imaging studies that characterize the dynamics of the processes and how they are disrupted by the absence of postsynaptic dynein. It will be important to determine when dynein is initially localized to the NMJ and whether dynein’s postsynaptic accumulation precedes the establishment of newly formed glutamate receptor clusters. Our studies further suggest novel roles for dynein in regulating PIP_2_ production and signaling. The deconstruction of the signaling cascades and physical interactions that regulate the postsynaptic cytoskeleton, synaptic terminal growth, and receptor organization are important challenges to understanding the mechanisms involved in synaptic function, plasticity, and related disease of the nervous system.

## Materials and Methods

### Drosophila lines

Flies were raised on standard food, and all crosses were performed at 29°C. *w^1118^* was used as the wildtype outcross control. Stocks used from the Bloomington Drosophila Stock Center are as follows: *UAS-dhc64cRNAi^HMS01587^*, *UAS-swRNAi^HM05249^*, *UAS-BSpecRNAi^GL01174^*, *Mef2-Gal4.R*-3, *UAS-sktlRNAi^JF02796^*, *UAS-PLCdelta-PH-EGFP-3*, *mhc-GluRIIA-Myc-*2, *mhc-GluRIIB-Myc*-3. Stocks used from Vienna Drosophila Resource Center (VDRC) are as follows: *UAS-khcRNAi^GD12278^* (44337). Other stocks used in this study: *G14-Gal4/CyoGFP* (Saitoe et al., 2001), *Dhc64c::Dhc64c^3xHA-WT^* and *Dhc64c::Dhc64c^3xHA-P1^* (Silvanovich et al., 2003), *UASp-6xMyc-sktl* (Jouette et al., 2019; S. Claret) elavGal80 ((Yang et al., 2009), *UAS-Nlg1-GFP* (Banovic et al., 2010; H. Aberle), *C57-Gal4*, *UAS-DlgS97-GFP* (Bachmann et al., 2004), *M12-Gal4*, also known as *5053A-Gal4* (Inaki et al., 2010). The genotype analyzed for each figure is provided in Supplemental Table 1. Also noted in this table, if the figure is an image, is the image type shown (single section, sum of sections, or a max projection of sections).

### Immunostaining

Wandering 3^rd^ instar larvae were dissected in HL3 saline (70mM NaCl, 5mM KCl, 20mM MgCl_2_, 10mM NaHCO_3_, 5mM trehalose, 11 mM sucrose, and 5mM Hepes, pH 7.2) by filleting the larva open in Sylgard coated dishes. Animals were fixed in 4% paraformaldehyde for 20 minutes for all antibodies except for GluRIIA. For GluRIIA antibody staining larvae were fixed in Bouin’s solution (Sigma-Aldrich, St. Louis, MO) for 5 minutes. Antibodies were used at the following concentrations: rat anti-HA 3F10 at 1:400 (Sigma-Aldrich), mouse anti-Myc 9B11 at 1:2000-1:3000 (Cell Signaling Technology, Danvers, MA), mouse anti acetylated α-tubulin 6-11B-1 Alexa488 conjugated 1:50 (Santa Cruz Biotechnology, Inc., Dallas, TX), mouse anti-Dhc64c P1H4 at 1:400 (McGrail and Hays, 1997), mouse anti-Dlg 4F3-c at 1:500 (Developmental Studies Hybridoma Bank (DSHB), Iowa City, IA), mouse anti-GluRIIA 8B4D2-s at 1:10 (DSHB), rabbit anti-GluRIIC at 1:3000 (Ramos et al., 2015; M. Serpe), rabbit anti-GluRIIB at 1:1000 (Ramos et al., 2015; M. Serpe), rabbit anti-sktl 1:800 (Claret et al., 2014; S. Claret), mouse anti-Scar P1C1 at 1:50 (DSHB), mouse anti-α-Spectrin 3A9-s at 1:10 (DSHB), guinea pig anti-β-Spectrin at 1:1000 (this study), rabbit anti-GFP TP401 at 1:1000 (Torrey Pines Biolabs, Secaucus, NJ), mouse anti-BRP nc82-s at 1:25 (DSHB), mouse anti-Hts 1B1-c at 1:100 (DSHB), mouse anti-Ten-m mAb20-s at 1:10 (DSHB), rabbit anti-Kinesin Kin01 at 1:200 (Cytoskeleton, Inc., Denver, CO), rabbit anti Nlg1 1:500 (Banovic et al., 2010; S.Sigrist), guinea pig anti Nlg1 1:250 (Mozer and Sandstrom, 2012; B. Mozer). Secondary antibodies conjugated with Alexa488 or Alexa594 were used at 1:500 (Jackson ImmunoResearch Laboratories, Inc., West Grove, PA). Samples were mounted in Prolong Glass antifade mounting media (Thermo Fisher Scientific, Waltham, MA).

### Bouton number per muscle area quantification

Wandering 3^rd^ instar larvae from crosses of Mef2Gal4 to W^1118^ or UAS-dhc64cRNAi^HMS01587^ were dissected and processed as described in the immunostaining methods section. Images were acquired on a Zeiss Cell Observer Spinning disk confocal microscope using a Photometrics QuantEM 512SC camera and laser lines at 488nm and 561nm and emitted fluorescence was collected through 503-538nm and 580-653nm emission filters. Zen2 software was used to acquire the images. A 10x EC Plan NeoFluor 0.3NA objective (1.33μm pixel size) was used to image muscles for area quantifications. A 63x Plan Apo 1.4NA objective (0.21μm pixel size) was used to image the NMJ at abdominal segment A2 of muscles 6/7 and abdominal segment A3 of muscle 4. When necessary, tiling of the NMJs was done and images were stitched together using the Zen software. Ten female animals of each genotype were dissected and imaged. The total muscle areas were obtained by tracing the outline of the muscle in Fiji. Boutons were identified by eye and counted for NMJ6/7 and 1b boutons were counted for NMJ4.

### Quantification of protein intensities at neuromuscular junctions

Wandering 3^rd^ instar larvae from crosses were dissected and processed as described in the immunostaining methods section. 6 female animals for each genotype were used. Animals were stained with the same primary and secondary antibody solutions and imaged under the same conditions. Samples were imaged in a Zeiss Cell Observer Spinning disk confocal microscope with a Yokogawa CSU-X1 spinning disk confocal head. Illumination was provided with lasers of 488 and 561 nm and emitted fluorescence was collected through 503-538, and 580-653 nm emission filters, respectively. Zen2 software (Zeiss), a QuantEM 512SC camera (Photometrics), and a 63x Plan Apo 1.4NA objective (0.21μm pixel size), with a Z-step size of 0.24μm were used to acquire the images. NMJ images were acquired at abdominal segments A3 and A4 of muscle 4. For analysis the last bouton of a synaptic terminal was selected. A Jython script was written for Fiji to measure the postsynaptic protein intensities at NMJs. Briefly, a 47 x 47 pixel area (9.96um X9.96um) was selected from the maximum intensity projection. The selected area was used to crop all slices in the original image, creating a 3D stack. Each channel underwent background correction by subtracting the average of the signal below the Otsu threshold from the original image. To estimate the location of the membrane region, we first thresholded the original image using the Otsu algorithm. The mask was then cleaned up using a morphological opening operation. Finally, the eroded mask was subtracted from a dilated version of the mask, selecting a 3D shell around the boundary of the cleaned-up mask. All morphological operations were conducted in 3D and with a structuring element of a sphere with radius 2 pixels, or 0.42 um. Within the shell, the average intensity for each channel was measured. For the Dlg and β-Spectrin intensities, Dlg was measured as *HRP-CH 1* and β-Spectrin was measured as *CH 2*. All measurements were performed in the background-corrected cropped image. The code is available on the GitHub repository https://github.com/tp81/neisch-dynein-2021.

### Quantification of glutamate receptor field sizes

Wandering 3^rd^ instar larvae from crosses were dissected and processed as described in the immunostaining methods section. 6 female animals for each genotype were used. Animals were stained with the same primary and secondary antibody solutions and imaged under the same conditions. Samples were imaged in a Zeiss Cell Observer Spinning disk confocal microscope (Zeiss, Obekochen, Germany) with a Yokogawa CSU-X1 spinning disk confocal head (Yokogawa, Tokyo, Japan). Illumination was provided with lasers of 488 and 561 nm and emitted fluorescence was collected through 503-538, and 580-653 nm emission filters, respectively. Zen2 software (Zeiss), a QuantEM 512SC camera (Photometrics, Tuscon, AZ), and a 63x Plan Apo 1.4NA objective (0.21μm pixel size), with a Z-step size of 0.24μm were used to acquire the images. NMJ images were acquired at abdominal segments A3 and A4 of muscle 4. To quantify the receptor field size, a script was written in Jython within Fiji. The HRP channel *CH 1* was thresholded using the “Moments” algorithm to each slice. Each slice of the glutamate receptor subunit channel *CH 2* underwent rolling ball background subtraction with radius 2 pixels (0.42um). A 3D mask was created by applying the Bersen local thresholding algorithm with radius 10 (2.1 um) to each slice. All parts in the second mask that were not included in the first were excluded from further analysis. The 3D object counter was used to count and measure all objects with a volume greater than 10 voxels (0.10584 μm^3^). The volume was used to estimate an equivalent sphere radius by using the following formula: r_eq_ = ∛ ( V / (4/3 *×* π)). All code is available on the GitHub repository https://github.com/tp81/neisch-dynein-2021.

### Quantification of BRP punctae per glutamate receptor field

Reconstructed 3D Structural Illumination Microscopy (SIM) images were analyzed using Imaris 9.5.1 (Bitplane, Belfast, UK). The region of interest containing NMJ4 was first selected. To quantify BRP punctae per image, the BRP channel was defined as spots. For the algorithm settings, different spot sizes (region growing) was enabled, as was the shortest distance calculation. The estimated XY-diameter was 0.2μm and the estimated Z-diameter was 0.8μm. Background subtraction was enabled. Region growing used local contrast and was manually thresholded. Spots were classified by filtering by quality and manually adjusted for spots visually observed as true signal in the image. To quantify the GluRIIC receptor fields per image, the GluRIIC channel was defined as surfaces. For the algorithm settings, the shortest distance calculation was enabled. Smoothing was applied with a surface detail of 0.065μm. The background was subtracted with a diameter of the largest sphere which fits into the object being 0.244μm. Thresholding was manually defined to match the visible signal. Split touching objects (region growing) was enabled with a seed point diameter of 0.6μm. Seed points were classified by a quality filter, and manually adjusted. The total number of BRP punctae observed in each NMJ4 image analyzed was divided by the total number of GluRIIC receptor fields observed in the same image to determine the BRP/GluRIIC ratio.

### Electrophysiology

To measure mEJPs, wandering 3^rd^ instar female larvae from crosses of *elavGal80/Y; Mef2Gal4* to *w^1118^* or *UAS-dhc64cRNAi^HMS01587^* were dissected in HL3 saline and visceral organs were removed. Segmental nerves were severed near the ventral nerve cord and the brain was then fully removed. The HL3 saline in the filleted larval prep was replaced HL3 saline containing 1.5mM CaCl_2_ for mEJPs recordings. Recordings were performed on muscle 6. of abdominal segments A2 and A3 using sharp glass microelectrodes filled with 3M KCl. Only recordings with a resting membrane potential of −60 mV or lower were analyzed. Recordings were performed over multiple days. Measurement of mEJPs were performed using Mini Analysis (Synaptosoft, Inc., Decatur, GA). The average recording time for controls was 1.8 minutes and for dynein depletion the average recording time was 1.7 minutes.

### Structured Illumination Microscopy (SIM)

Images were acquired in a Nikon Ti-E inverted microscope (Nikon Inc., Melville, NY) equipped with a CFI SR Apo 100x TIRF oil immersion objective lens, NA 1.49 (0.033 μm pixel size). Illumination was provided by 488 and 561 nm lasers fed through a multiple mode fiber to a Nikon N-SIM illuminator with the appropriate diffraction grating. The cognate emission filters used were 522-545 and 605-670 nm, respectively. Images were acquired in 3D SIM mode with an Orca-Flash4.0 V3 sCMOS camera and the SIM image reconstructed using Nikon software and the default software settings for illumination modulation contrast, high resolution noise suppression, and out of focus blur suppression (Nikon Elements 5.11). Pixel size of the reconstructed image is 33 nm.

### β-Spectrin antibody production

A DNA sequence encoding *Drosophila* β-Spectrin amino acids 1013-1463 was PCR amplified from pUASTattB-β-Spectrin (Avery et al., 2017) using the oligonucleotides: 5’GGAATTCCATATGGAGCGCGAAGCCAACAGCATC-3’ and 5’ACCGCTCGAGCGTCTTCTTCACCACAATCGGTTCG-3’. The PCR product was digested and with Nde1/Xho1 restriction enzymes and inserted into the bacterial expression vector pET-30a(+) (Novagen, Madison, WI) cut with the same enzymes, to introduce a His tag at the C-terminus of the β-Spectrin protein fragment. The resulting construct was confirmed by DNA sequencing. The β-Spectrin-His construct was transformed into BL21(DE3) cells (Novagen) and induced for protein expression with 0.5mM IPTG (Invitrogen, Carlsbad, CA). The bacterial pellet was lysed by sonication in binding buffer consisting of PBS with 10mM imidazole (Sigma-Aldrich) and Complete Protease Inhibitor Cocktail, EDTA-Free (Roche, Mannheim, Germany). After clarification by centrifugation, the lysate was incubated with Ni-NTA agarose resin for binding (Invitrogen), and later the resin was transferred to a Poly-Prep chromatography column (BioRad, Hercules, CA). The β-Spectrin-His protein was eluted in PBS containing 200 mM imidazole, and later dialyzed in PBS to remove imidazole. β-Spectrin-His protein was injected into a Guinea Pig at Pocono Rabbit Farm and Laboratory, Inc. (Canadensis, PA), and serum containing β-Spectrin antibody collected.

### Dlic-GFP generation

A PCR fragment encoding eGFP was first ligated to the 3’ end of dlic coding sequences. The Dlic-GFP fragment was then fused to ∼2kb genomic sequences including 5’ UTR and upstream regulated region used for the dlic genomic transgene (Mische et al., 2008). The final construct was cloned into the pCaspeR4 vector and transgenic flies were generated. The Dlic-GFP transgene is fully functional for rescuing to viability dlic lethal mutations.

## Acknowledgements

We thank Sandra Claret, Brian Mozer, Ela Serpe, Stephan Sigrist, and the Developmental Studies Hybridoma Bank (DSHB) for providing antibodies; Hermann Aberle, Sandra Claret, and Claude Desplan for providing fly lines; Myung-Jun Kim for training and assistance with electrophysiology experiments and analysis; Guillermo Marqués for assistance with SIM imaging; and Akshaya Gupta for technical assistance with the β-Spectrin antibody production. Thanks to Hiroshi Nakato, Mike O’Connor, Aidan Peterson and Guillermo Marqués for helpful comments and suggestions on the manuscript. Image acquisition was done at the University Imaging Centers at the University of Minnesota.

---

**Figure S1. Dynein’s localization at the postsynaptic NMJ is not dependent on Kinesin-1, which predominantly localizes to the presynaptic side of the NMJ.**

**A-A’)** The dynein light intermediate chain subunit, dlic, tagged with GFP and expressed under its endogenous promoter, has a punctate postsynaptic localization. This localization is similar to that observed for the dynein heavy chain subunit, dhc64c, using an antibody that is specific. **B-B’)** Kinesin-1, tagged with GFP and expressed ubiquitously under the α-tubulin promoter, has a strong presynaptic localization at the NMJ, and no specific postsynaptic localization. **C-C’)** The localization of dynein at the NMJ in control animals. **D-D’)** The localization of dynein at the NMJ is not affected when kinesin-1 is depleted from the muscle by RNAi. Halos of dynein staining are observed around nuclei, labeled with the letter N, when kinesin-1 is depleted from the muscle, which is not observed in controls. Boxed areas are shown as insets. In all images, HRP labels the presynaptic side of the NMJ. Scale, 10μm.

**Figure S2. Dynein is required in the muscle for synaptic growth and affects the size of glutamate receptor clusters at the NMJ.**

To verify that the depletion of dynein specifically within the muscle affects synaptic growth of the NMJ, the number of boutons and muscle area were compared between control animals, animals depleted of muscle dynein using the *Mef2-Gal4* driver, and animals depleted of dynein using the *Mef2-Gal4* driver co-expressing *elav-Gal80* to suppress any possible neuronal dynein depletion. The number of boutons and total muscle area was measured for muscle 4. Measurement for control animals, and dynein depletion by *Mef2-Gal4* are values taken from Figure 2. **A)** The number of boutons per muscle area at NMJ4 is significantly reduced when dynein is depleted in the muscle and any nervous system expression is suppressed with *elav-Gal80*. **A’)** Similar to what was observed for depletion of dynein under *Mef2-Gal4* alone, the muscle area is not significantly affected when *elav-Gal80* is co-expressed. n=number of NMJs analyzed. Unpaired two-tailed t-test results: **, P<0.005, ns=not significant. **B)** Intensity values of GluRIIA, GluRIIB, and GluRIIC at the NMJ in control animals, and those depleted of dynein by RNAi. n=number of NMJ analyzed. Unpaired, two-tailed t-test results: ns=not significant. **C-C’)** A representative image of GluRIIA localization at the NMJ is controls and when dynein is depleted in the muscle **(D-D’)**. **E-E’)** A representative image of GluRIIB localization at the NMJ is controls and when dynein is depleted in the muscle **(F-F’)**. **G-G’)** Myc-GluRIIA localization at the NMJ in a control animal and in an animal depleted of dynein in the muscle **(H-H’)**. **I-I’)** Myc-GluRIIB localization at the NMJ in a control animal and in an animal depleted of dynein in the muscle **(J-J’)**. In all images, HRP labels the presynaptic side of the NMJ. Scale, 10μm.

**Figure S3. A β-Spectrin antibody shows specificity.**

In order to visualize β-Spectrin at the NMJ, a new antibody was generated. We tested the specificity of this antibody using muscle specific depletion of β-Spectrin. **A-A’’)** In control animals β-Spectrin is enriched on the postsynaptic side of the NMJ. **B-B’’)** When β-Spectrin is depleted from the muscle, presynaptic β-Spectrin is now able to be visualized with the antibody, as indicated by the arrows. Black arrowheads indicate trachea that stain positive for β-Spectrin. HRP labels the presynaptic side of the NMJ. Scale, 10μm.

**Figure S4. Dynein depletion in the muscle does not affect the postsynaptic localization of the transmembrane proteins Ten-m or Nlg1.**

**A-A’)** The localization of Nlg1-GFP at the NMJ4 in a control animal. **B-B’)** Depletion of dynein from the muscle does not affect the localization of Nlg-GFP at the NMJ. **C-C’)** The wild type localization of Ten-m at NMJ4. **D-D’)** Depletion of dynein from the muscle does not affect to localization of Ten-m. **E-E’)** The localization of Ten-m at NMJ3 of control animals. **F-F’)** Ten-m localization at NMJ3 of a postsynaptic dynein depleted animal. **G)** quantification of postsynaptic Ten-m intensity levels in control and dynein depleted animals. n=number of NMJs analyzed. Unpaired two-tailed t-test results: ns=not significant. **H-H’)** The localization of Nlg1 at NMJ4 using a guinea Pig anti-Nlg1 antibody in a control animal and in an animal depleted of postsynaptic dynein **(I-I’)**. **J-J’)** The localization of Nlg1 at NMJ4 using a rabbit anti-Nlg1 antibody in a control animal and in an animal depleted of postsynaptic dynein **(K-K’)**. Boxed areas are shown as insets. In all images, HRP labels the presynaptic side of the NMJ. Scale, 10μm.

**Figure S5. Sktl can be effectively depleted from the postsynaptic muscle using RNAi.**

Sktl was strongly depleted from muscle 12 (left side of the dotted lines), by expressing *sktlRNAi* with the *M12-Gal4* driver **(A,B)**. Red arrowheads indicate the position of muscle 12 NMJs. Muscle 13 (right side of the dotted lines) which does not express *sktlRNAi* is shown for comparison. Black arrowheads indicate the position of muscle 13 NMJs. To visualize Sktl, a Sktl specific antibody was used. HRP labels the presynaptic side of the NMJ **(A’,B’)**. **C-D’)** The maximum of the Look Up Table (LUT) for images visualizing the localization of GluRIIC in control and Sktl depleted animals shown in Figure 8A,B was adjusted to readily discern the extra synaptic punctae noted in controls that were decreased in Sktl depleted animals. Both images were adjusted with the same LUT maximums. Scale, 10μm.

## Notes

### Competing Interest Statement

The authors have declared no competing interest.

